# Partial loss of CFIm25 causes aberrant alternative polyadenylation and learning deficits

**DOI:** 10.1101/735597

**Authors:** Callison E. Alcott, Hari Krishna Yalamanchili, Ping Ji, Meike E. van der Heijden, Alexander B. Saltzman, Mei Leng, Bhoomi Bhatt, Shuang Hao, Qi Wang, Afaf Saliba, Jianrong Tang, Anna Malovannaya, Eric J. Wagner, Zhandong Liu, Huda Y. Zoghbi

## Abstract

We previously showed that *NUDT21*-spanning copy-number variations (CNVs) are associated with intellectual disability (Gennarino et al., 2015). However, the patients’ CNVs also included other genes. To determine if reduced *NUDT21* function alone can cause disease, we generated *Nudt21*^+/-^ mice to mimic the human state of decreased expression. We found that although these mice have 50% reduced *Nudt21* mRNA, they only have 30% less of its cognate protein, CFIm25. Despite this partial protein-level compensation, the *Nudt21*^+/-^ mice have learning deficits and cortical hyperexcitability. Further, to determine the molecular mechanism driving neural dysfunction, we partially inhibited *NUDT21* in human stem cell-derived neurons to reduce CFIm25 by 30%. This reduction in CFIm25 was sufficient to induce misregulated alternative polyadenylation (APA) and protein levels in hundreds of genes, dozens of which cause intellectual disability when mutated. Altogether, these results indicate that disruption of *NUDT21*-regulated APA events in the brain can cause intellectual disability.

## INTRODUCTION

The brain is acutely sensitive to the dose of numerous proteins, such that even small perturbations in their levels can cause neurological disease. Proteins that affect the expression of other genes, such as transcriptional or translational regulators, are particularly critical (Rubeis et al., 2014; Vissers et al., 2015; Yin and Schaaf, 2017). Canonical examples of such broad regulators are the RNA-binding protein FMR1, which underlies fragile X syndrome, and the chromatin modulator MeCP2, whose loss or gain respectively causes Rett syndrome or *MECP2*-duplication syndrome (Amir et al., 1999; Pieretti et al., 1991; Verkerk et al., 1991).

Alternative polyadenylation (APA) is an important mechanism of transcript and protein-level regulation. Indeed, nearly 70% of human genes have multiple polyadenylation [poly(A)] sites, usually within the 3′ untranslated region (UTR) of the mRNA, allowing for transcript isoforms with different 3′ UTR lengths and content (Derti et al., 2012; Hoque et al., 2013). The general implication is that longer 3′ UTRs contain additional regulatory motifs, such as miRNA and RNA-binding protein binding sites. This allows for differential regulation of those gene products, typically through reduced mRNA stability, but also through other mRNA metabolism mechanisms, such as translation efficiency and localization [reviewed in (Tian and Manley, 2016)]. Alternative poly(A) site usage has been observed in multiple biological contexts, both in different disease states and in normal physiology and development (Hoque et al., 2013; Masamha and Wagner, 2017). In general, as cells differentiate, pathway-specific mRNAs increasingly use distal polyadenylation sites, while cells that are more proliferative use more proximal poly(A) sites (Hoque et al., 2013; Ji et al., 2009; Sandberg et al., 2008). In the brain, average 3’ UTR length increases throughout neurogenesis, and mature, post-mitotic neurons have the longest average 3’ UTR length of any cell type (Guvenek and Tian, 2018; Miura et al., 2013; Zhang et al., 2005).

Numerous genes regulate alternative polyadenylation, but *NUDT21* is among the most consequential (Gruber et al., 2012; Masamha et al., 2014; Tian and Manley, 2016). *NUDT21* encodes CFIm25, a component of the mammalian cleavage factor I (CFIm) complex (Kim et al., 2010; Rüegsegger et al., 1996; Yang et al., 2011). CFIm25 binds UGUA sequences in pre-mRNA and the CFIm complex helps recruit the enzymes required for cleavage and polyadenylation (Brown and Gilmartin, 2003; Rüegsegger et al., 1998; Yang et al., 2011, 2010; Zhu et al., 2018). The UGUA binding sites are often enriched at the distal polyadenylation sites of *NUDT21*-regulated RNAs, so CFIm25 typically promotes the synthesis of longer mRNA isoforms (Zhu et al., 2018). Thus, when *NUDT21* expression is reduced, proximal cleavage sites are more frequently used. Indeed, CFIm25 downregulation in multiple human and mouse cell lines causes 3′ UTR shortening in hundreds of genes, and a consequent increase in protein levels of a subset of those genes (Brumbaugh et al., 2018; Gennarino et al., 2015; Gruber et al., 2012; Kubo et al., 2006; Li et al., 2015; Martin et al., 2012; Masamha et al., 2014). Notably, *MECP2* is among the most affected genes in these cell-line studies, and slight perturbations in MeCP2 levels cause neurological disease (Chao and Zoghbi, 2012). Moreover, *NUDT21* is a highly constrained gene. In the Genome Aggregation Database (gnomAD) of ∼140,000 putatively healthy individuals, 125 missense and 13 loss of function variants would be expected in *NUDT21* if loss of function were not pathogenic, but instead there are only 15 missense and zero loss-of-function variants, suggesting that *NUDT21* loss of function is incompatible with health (Lek et al., 2016). Given this evidence, we hypothesized that *NUDT21* variants can cause neurological disease through APA misregulation of *MECP2* and other dose-sensitive genes in neurons.

Combining results from our previous work with data from the Decipher database, we have identified nine individuals with *NUDT21*-spanning duplications that have intellectual disability, and two patients with deletions that have both intellectual disability and seizures (Firth et al., 2009; Gennarino et al., 2015). However, the duplication patients also have three other genes common to their copy-number variations (CNVs) and the deletion patients have nearly 20 common genes (Gennarino et al., 2015). CNVs of these other genes could be causing their symptoms. Therefore, it is important to determine if *NUDT21* loss of function alone is sufficient to cause disease. Identifying the disease-causing genes within CNVs facilitates more accurate diagnosis and prognosis, and allows for targeted therapy development. To that end, we generated *Nudt21*^+/-^ mice to model the reduced CFIm25 expression observed in humans and assessed for phenotypes similar to the symptoms seen in the deletion patients. We found that *Nudt21* heterozygosity causes a 50% loss of wild-type *Nudt21* mRNA as expected, but only a 30% reduction of its cognate protein, CFIm25. Consistent with what we observed in the deletion patients, we found that *Nudt21* loss of function is sufficient to cause learning deficits in mice using a variety of behavioral assays. Further, to see how *NUDT21* loss might lead to disease, we analyzed *NUDT21*-depleted human neurons for aberrant alternative polyadenylation with 3’ end sequencing and for protein level aberrations with quantitative mass spectrometry. These approaches showed that a 30% reduction of CFIm25 in human neurons induces widespread abnormal alternative polyadenylation and protein levels, including for dozens of dose-sensitive, disease-associated genes. Altogether, these results provide important in vivo and human-specific evidence that reduced *NUDT21* expression can lead to intellectual disability.

## RESULTS

### *Nudt21* heterozygotes have 50% less *Nudt21* mRNA in their brain, but only 30% reduced CFIm25 protein

*Nudt21* expression has not been well explored within organisms, particularly in the brain. Thus, we first sought to confirm that mice express *Nudt21* in the brain, and found that we could readily detect CFIm25 in neuronal nuclei in regions important for learning and memory—the cortex and hippocampus (***Figure 1A***). We then generated *Nudt21* knockout mice by excising exons two and three, which removes the RNA-binding domain of CFIm25 and induces a frame shift mutation leading to nonsense-mediated decay of the transcript (***Figure 1B***) (Yang et al., 2011). We crossed the *Nudt21*^+/-^ mice with each other and observed that of the 32 pups born, there were no homozygous null offspring, indicating that *Nudt21* is essential for life (***Figure 1C***). Unexpectedly, while the heterozygous mice have the anticipated 50% reduction in wildtype *Nudt21* mRNA in whole-brain extracts, there was only ∼30% reduction in CFIm25 protein (***Figures 1D& E***). These results demonstrate the successful knockout of one *Nudt21* copy and reveal a partial post-transcriptional compensation of CFIm25 protein levels. Finally, we found that the *Nudt21*^+/-^ mice had a normal life span, but consistently weighed ∼10% less than their wild-type littermates, indicating that *Nudt21* heterozygosity might cause some abnormalities (***Figure 1—figure supplement 1***).

**Figure 1.**
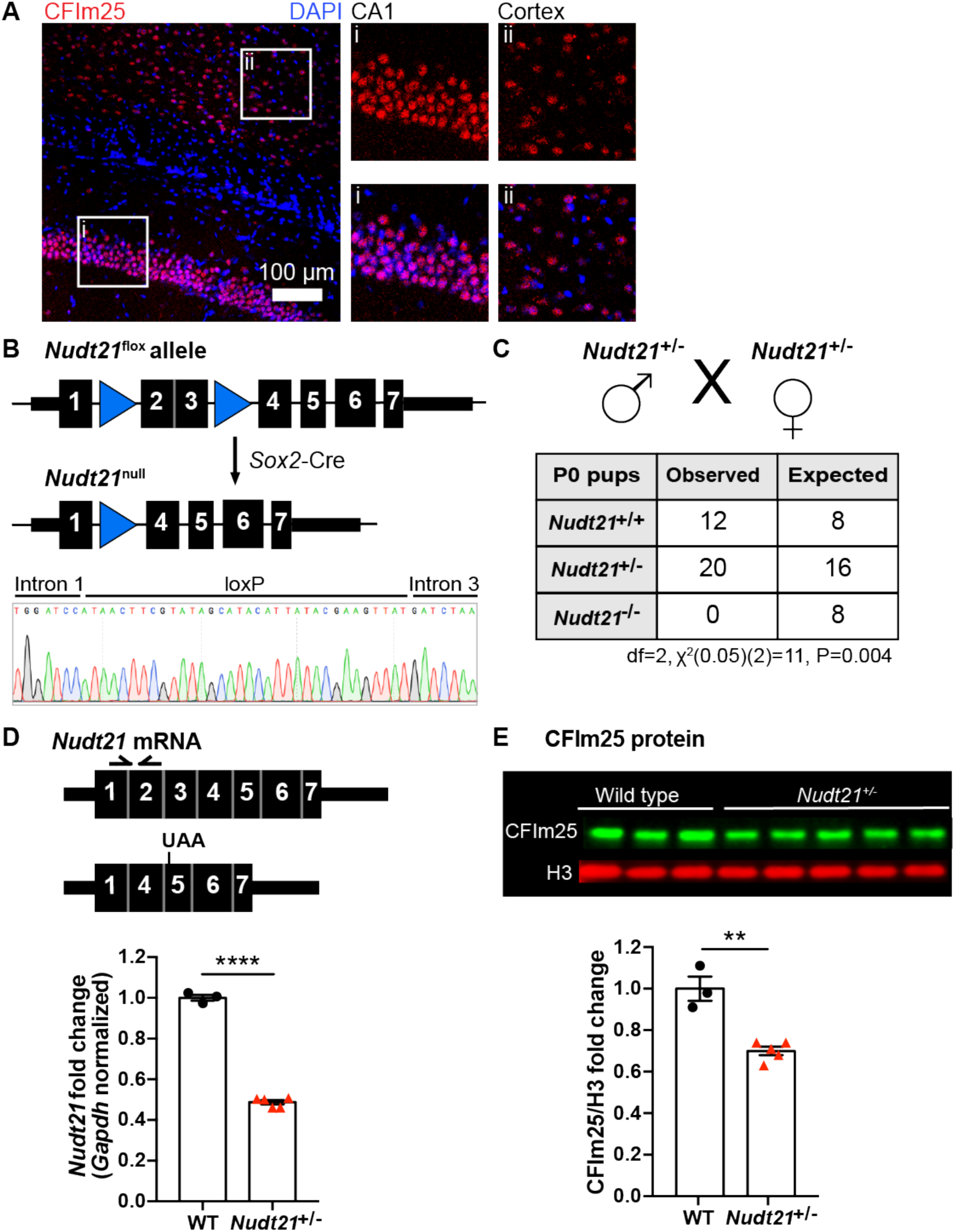
*Nudt21* heterozygotes have 50% less *Nudt21* mRNA in their brains, but only 30% reduced CFIm25 protein. **(A)** Immunofluorescence showing CFIm25 expression in the (i) mouse hippocampus and (ii) cortex. **(B)** Schematic of the floxed and recombined *Nudt21* alleles (top), and sequencing showing successful recombination (bottom). **(C)** Observed and expected offspring counts born with each possible genotype from *Nudt21*^+/-^ mating pairs. No homozygous *Nudt21* null offspring were born, when eight would be expected if loss of *Nudt21* did not affect survival: P=0.004, analyzed by two-tailed, chi-square test. **(D)** Schematic of wild-type and recombined *Nudt21* mRNA (top). Arrows indicate RT-qPCR primer binding sites, and UAA shows site of induced premature stop codon after recombination. RT-qPCR analysis shows expected 50% reduction of whole-brain, *Gapdh*-normalized, wild-type *Nudt21* mRNA in five-week-old mice with one wild-type *Nudt21* allele and one recombined, null allele (bottom): P<0.0001, n=3-5/genotype. **(E)** Western blot image comparing five-week-old *Nudt21*^+/-^ mice CFIm25 protein levels with their WT littermates (top). Western blot analysis showing ∼30% reduction of H3-normalized CFIm25 protein levels in *Nudt21*^+/-^ mice: P=0.0012, n=3-5/genotype. We confirmed that CFIm25 does not regulate H3. For all charts, error bars indicate SEM. All data analyzed by unpaired, two-tailed t-test unless otherwise stated. **P<0.01; ****P<0.0001. Weights of the heterozygous animals are shown in ***Figure 1—figure supplement 1***.

**Figure 1—figure supplement 1.**
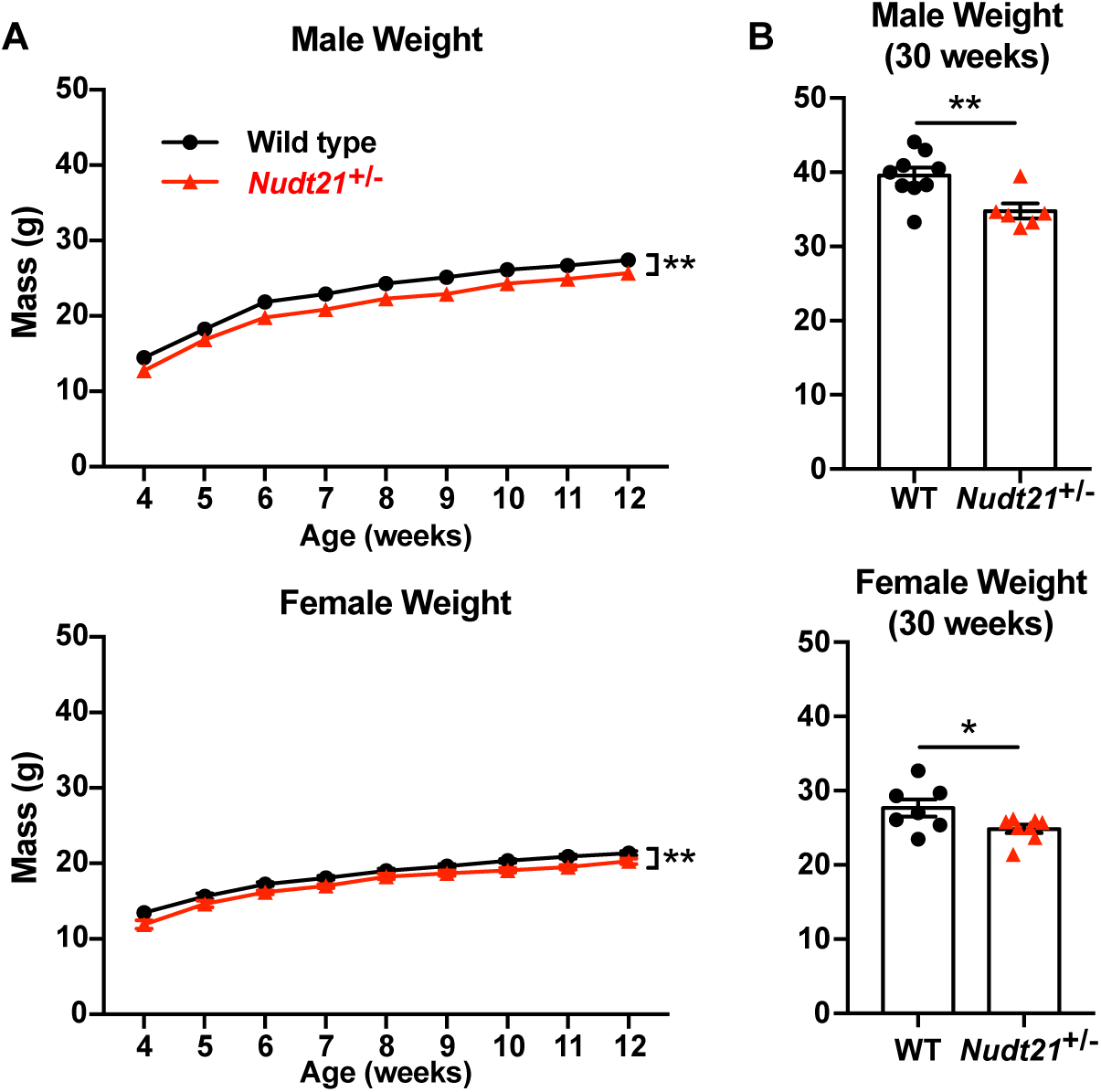
*Nudt21*^+/-^ mice weigh less. **(A)** Male (top) and female (bottom) *Nudt21*^+/-^ mice weigh less than their wild-type littermates from weening to twelve weeks: P=0.002 (males) and P=0.004 (females), analyzed by two-way repeated measures ANOVA (genotype*week). n=15-26/genotype. **(B)** Reduced weight persists up to at least 30 weeks in male (top) and female (bottom) *Nudt21*^+/-^ mice: P=0.008 (males) and P=0.04 (females), analyzed by unpaired, two-tailed t-test. n=6-9/genotype. Error bars indicate SEM. *P<0.05; **P<0.01.

### Partial loss of CFIm25 causes learning deficits

To determine if *Nudt21* heterozygosity affects cognitive function, we compared *Nud21*^+/-^ mice and their wild-type littermates in several neurobehavioral assays starting at 30 weeks of age. We focused on learning and memory assays because intellectual disability is the most pronounced and consistent symptom seen in patients with *NUDT21*-spanning CNVs (Gennarino et al., 2015). We started with the conditioned fear test, which assesses the mice’s ability to learn to associate an aversive event with a sensory context. In this test, we initially train the mice by exposing them to two tone-shock pairings in a novel chamber. Healthy mice will freeze as a behavioral expression of fear when they hear the tone for the second time. The following day, we return the mice to the same chamber, where healthy mice who learned to associate the chamber with the shock will freeze in recognition of the chamber. Lastly, we place the mice into a new chamber and replay the tone, where the healthy mice will remember that the tone is associated with the shock and freeze in fear. We found that *Nud21*^+/-^ mice show abnormal behavior throughout the assay. Their initial fear learning was attenuated: when exposed to the reinforcement sound cue played again two minutes after the initial tone-shock pairing during training, they froze on average ∼50% less often than their wild-type littermates (***Figure 2Ai***). Furthermore, the following day, they froze on average ∼30% less in both tests, indicating that *Nud21*^+/-^ mice have reduced memory of the chamber or the sound cue compared to their wild-type littermates (***Figure 2Aii& iii***).

**Figure 2.**
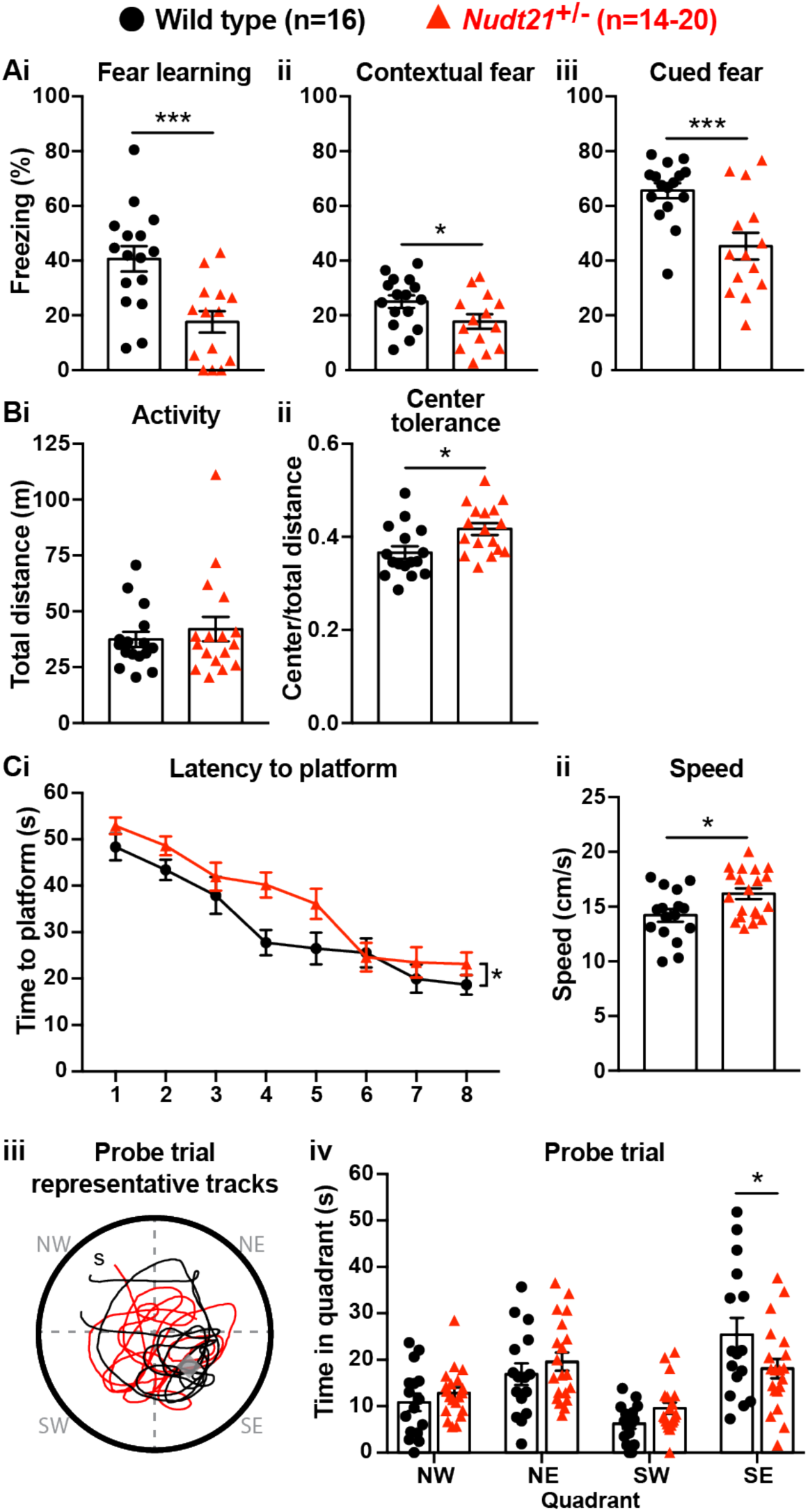
Partial loss of *Nudt21*-encoded CFIm25 protein causes learning deficits. **(A)** *Nudt21*^+/-^ mice have conditioned fear learning deficits: fear learning, P=0.0009; contextual fear, P=0.045; cued fear, P=0.0009. **(B)** The open field assay shows that *Nudt21*^+/-^ mice are no more active **(i)** but spend relatively more time in the center of the open field **(ii)**, indicating reduced anxiety: P=0.01. **(C)** *Nudt21*^+/-^ mice have spatial learning deficits in the Morris water maze. **(i)** They take longer to find the hidden platform during the training blocks (P=0.026, two-way, repeated measures ANOVA), **(ii)** despite swimming faster and farther (P=0.013). **(iii & iv)** When the hidden platform is removed in the probe trial, they spend less time in the quadrant that previously had the platform (P=0.039, Sidak’s multiple comparisons test). For all assays, mice were between 30-40 weeks of age. Error bars indicate SEM. Representative tracks are from the animal with the median result in each genotype. All data were analyzed by unpaired, two-tailed t-test unless otherwise stated. *P<0.05; ***P<0.001.

It is important to ensure that potential hyperactivity in the mutant mice does not confound the conditioned fear test results. If the mice are hyperactive, they may freeze less in the conditioned fear assay even if they remember the chamber or the cue. Therefore, we tested our mice in the open field assay. We place the mice in an open chamber and record the distance they travel in 30 minutes. We also record how much time they spend in the exposed center part of the chamber compared to the more hidden periphery. In this assay, hyperactive mice cover a greater distance, and anxious mice spend more time on the perimeter of the chamber (Crawley, 1985).

We found that the *Nud21*^+/-^ mice were not hyperactive, thus validating the conditioned fear results by showing they are not confounded by hyperactivity and indicating that partial loss of *Nudt21* function does indeed cause cognitive defects (***Figure 2Bi***). Intriguingly, the *Nud21*^+/-^ mice spent ∼15% more time in the exposed, center part of the open field, which suggests they have reduced anxiety and shows that they have neuropsychiatric features beyond learning and memory deficits (***Figure 2Bii***).

To determine if the learning deficits we detected in the *Nud21*^+/-^ mice extended to other learning paradigms, we tested spatial learning using the Morris water maze. In this assay, the mice are trained over four days to locate a hidden platform in a pool of water using visual cues provided on the perimeter of the room. We initially reveal the platform to the mice to confirm visual acuity. Then the mice are individually placed at different locations around the perimeter of the pool eight times per day and timed for how long it takes to locate the platform. Mice with spatial learning deficits require more time to find the platform throughout training. For an additional measurement, the platform is removed after the training and the mice are placed in a novel location on the perimeter of the pool. We then record how much time, out of a minute, they spend searching for the platform in the quadrant of the pool where it had been. Mice with spatial learning and memory deficits spend relatively less time in the platform quadrant, indicating that they do not remember its location as well. We found that the *Nudt21*^+/-^ mice indeed have spatial learning deficits. On average, they required more time to locate the platform during their training trials, despite swimming slightly faster and covering a greater distance (***Figure 2Ci& ii***). Moreover, when the platform was removed, the *Nudt21*^+/-^ mice spent ∼30% less time in the correct quadrant (***Figure 2Ciii& iv***). Thus, the conditioned fear and Morris water maze results show that *Nudt21*^+/-^ mice have learning and memory deficits in multiple domains.

### *Nudt21*^+/-^ mice have increased cerebral spike activity

In addition to intellectual disability, the patients with *NUDT21*-spanning deletions had seizures. In general, though, mice are much less sensitive to seizure-causing mutations than humans, but seizure susceptibility can be assessed using electroencephalography (EEG) (Amendola et al., 2014; Jiang et al., 1998; Kriscenski-Perry et al., 2002; Miura et al., 2002). Despite a lack of detectable seizures in *Nudt21*^+/-^ mice, we found significantly more EEG spikes in their frontal cortex relative to their wild-type littermates. This result indicates that *Nudt21* haploinsufficiency is alone sufficient to cause cortical hyperexcitability, and suggests that *NUDT21* loss of function might increase seizure risk (***Figure 3***).

**Figure 3.**
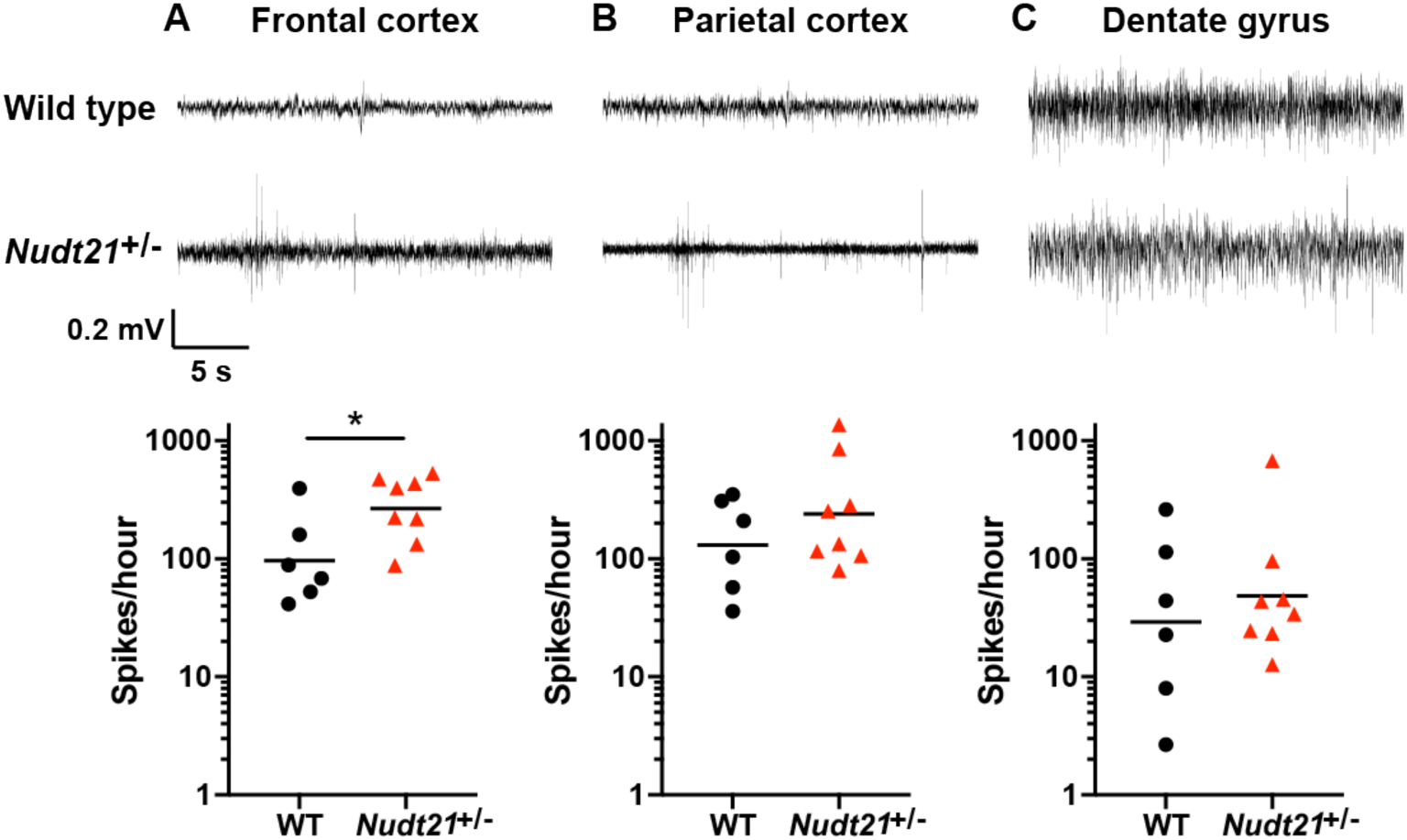
*Nudt21*^+/-^ mice have increased cerebral spike activity. **(A)** 57-week-old *Nudt21*^+/-^ mice have significantly increased spike activity in the frontal cortex by EEG (P=0.029), but not in the parietal cortex (**B)** and dentate gyrus (**C)**. Representative traces are on top and spike count summaries below. All data analyzed by two-tailed Mann-Whitney test. Central tendency lines show the geometric mean. *P<0.05.

### *NUDT21* depletion induces aberrant alternative polyadenylation and altered protein levels

The observations that individuals with *NUDT21*-spanning CNVs have intellectual disability (ID) and that *Nudt21*^+/-^ mice have learning deficits provides strong evidence that *NUDT21* loss of function causes disease, but do not reveal the molecular pathology. Therefore, to understand how partial loss of *NUDT21* function might sicken neurons and cause ID, we used shRNA to inhibit *NUDT21* in human embryonic stem cell (ESC)-derived glutamatergic neurons and assessed them for proteome and alternative polyadenylation (APA) dysregulation. To assess the proteome, we used mass spectrometry, and to directly measure APA events, we used poly(A)-ClickSeq (PAC-seq), an mRNA 3′-end sequencing method that allows for more accurate identification of polyadenylation sites than standard RNA sequencing (Elrod et al., 2019; Routh et al., 2017). The genotypes cluster by principle component in both assays, showing that both approaches are reproducible (***Figure 4—figure supplement 1A & B***).

**Figure 4.**
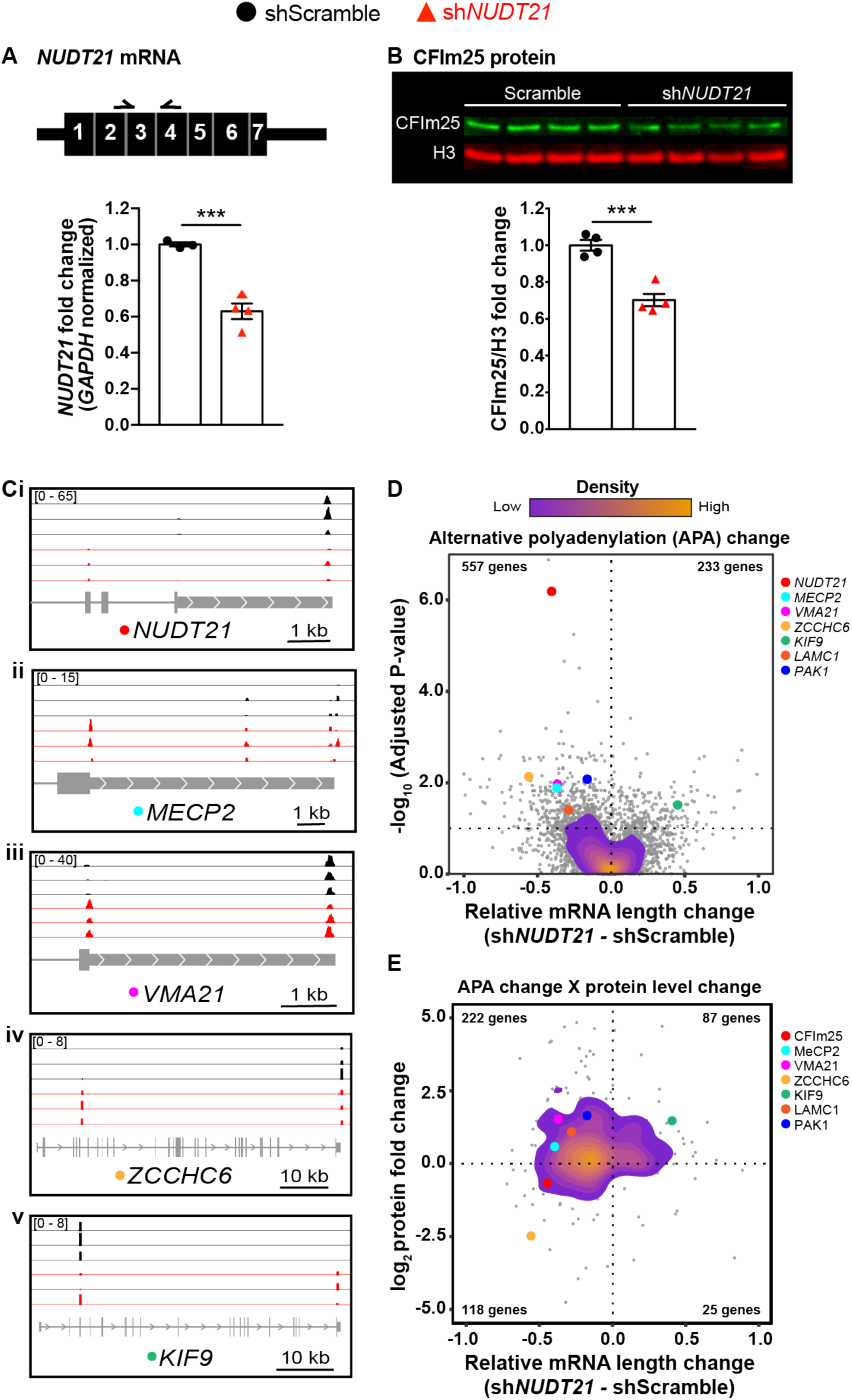
*NUDT21* depletion induces aberrant alternative polyadenylation and altered protein levels. **(A)** Schematic of *NUDT21* mRNA with qPCR primers (top) and RT-qPCR quantification of *NUDT21* mRNA levels in human embryonic stem cell (ESC)-derived neurons infected with scrambled shRNA (shScramble) or shRNA targeting *NUDT21* (sh*NUDT21*). The neurons infected with sh*NUDT21* have a 40% reduction of *GAPDH*-normalized *NUDT21*: P=0.0009, n=3-4/treatment. **(B)** Western blot image and quantification showing a 30% reduction of H3-normalized CFIm25 protein in sh*NUDT21*-infected neurons: P=0.0005, n=4/treatment. We confirmed that CFIm25 does not regulate H3. **(C)** Poly(A)-ClickSeq (PAC-seq) tracks showing reduced *NUDT21* and altered alternative polyadenylation in example genes: n=3/treatment. Peaks are 3′ end sequencing reads. Multiple peaks in a gene indicate multiple mRNA isoforms with different cleavage and polyadenylation sites. Bracketed numbers show counts per million; kb stands for kilobases. **(D)** Volcano plot showing relative mRNA length change in sh*NUDT21*-infected neurons compared to shScramble-infected controls. The horizontal, dashed line shows P_adjusted_=0.1, n=3/treatment. Principle component analysis shows sample clustering by treatment: (***Figure 4—figure supplement 1A***) **(E)** Mass spectrometry quantification of protein level fold change for genes with significantly altered APA (P_adjusted_ <0.1). For bar charts, error bars indicate SEM and data analyzed by unpaired, two-tailed t-test. ***P<0.001. Principle component analysis shows sample clustering by treatment: (***Figure 4— figure supplement 1B***). Source files for the PAC-seq and mass spectrometry quantification data are available in ***Figure 4—source data 1***.

**Figure 4—figure supplement 1.**
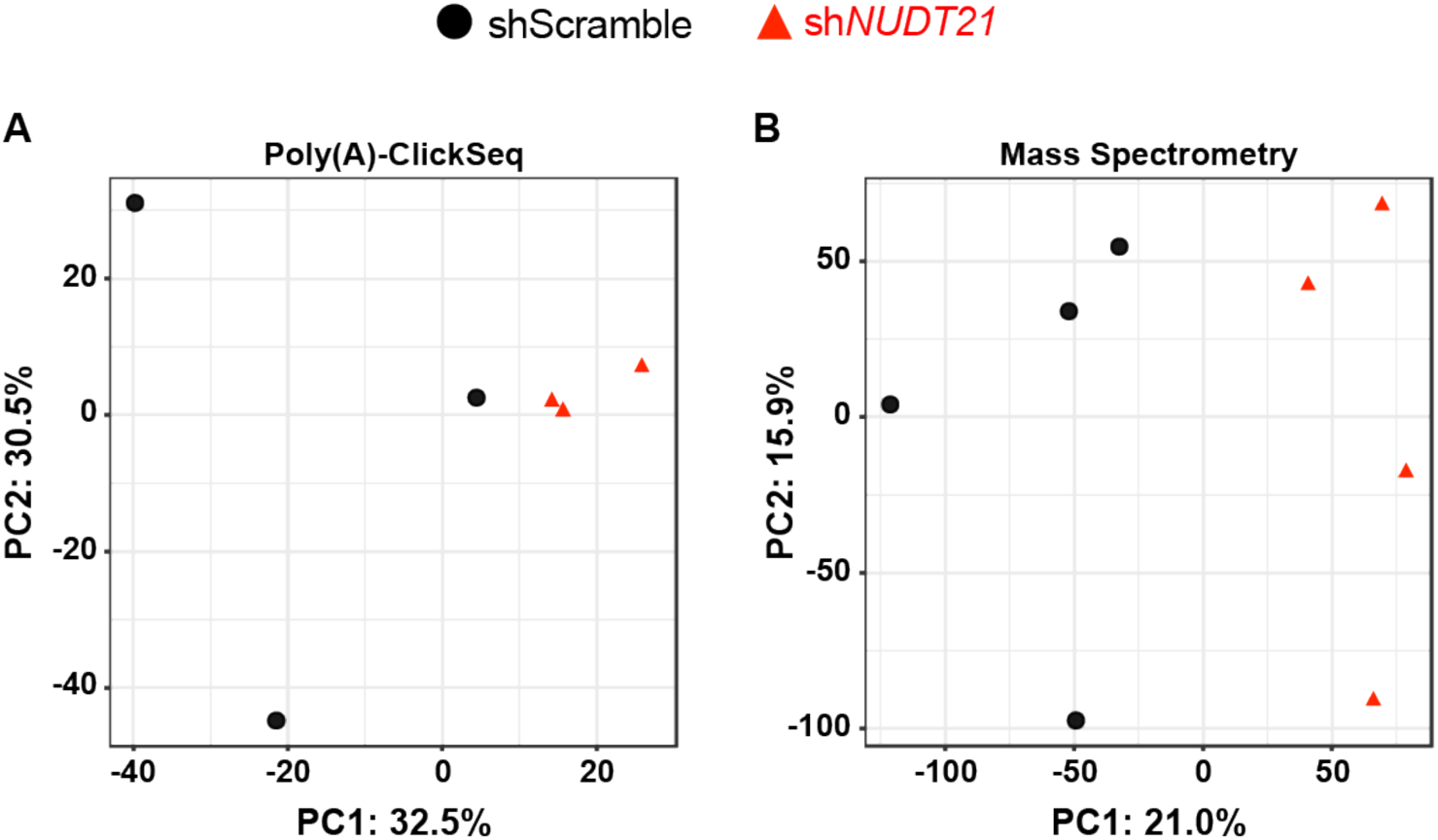
Control and sh*NUDT21*-infected human neurons transcriptomes and proteomes segregate by principle component analysis (PCA). **(A)** Control and sh*NUDT21*-infected human neurons segregate by principle component 1 (PC1) of differentially expressed genes. Principle components (%): 32.5, 30.5, 16.9, 11.2, 8.8, 0. **(B)** Control and sh*NUDT21*-infected human neurons segregate by PC1 of protein levels. Principle components (%): 21, 15.9, 15.3, 13.6, 12.4, 11.1, 10.6, 0. **Figure 4—source data 1. RNA length and protein level changes in genes with misregulated APA following neuronal *NUDT21* inhibition.** Likelihood ratio (lr), degrees of freedom (df), fold change (FC). **Supplemental data 1. Intellectual disability associations of genes with misregulated APA following neuronal *NUDT21* inhibition.** Probability of loss of function intolerance (pLI), intellectual disability (ID), Online Mendelian Inheritance in Man (OMIM), autosomal recessive (AR), autosomal dominant (AD), X-linked dominant (XLD), X-linked recessive (XLR)

Like in the whole-brain extracts from *Nudt21*^+/-^ mice, the human neurons had a 30% reduction of CFIm25 protein, despite a greater reduction of *NUDT21* mRNA (***Figure 4A & B***). This difference shows that there is also homeostatic stabilization of CFIm25 in human neurons. Intriguingly, in addition to reduced *NUDT21* mRNA levels as expected, the PAC-seq data also showed that *NUDT21* itself undergoes aberrant alternative polyadenylation following its depletion (***Figure 4Ci***). These shortened *NUDT21* mRNAs have no stop codon, and are likely to undergo non-stop decay, an mRNA decay mechanism that degrades mRNAs lacking an in-frame stop codon (Frischmeyer et al., 2002; van Hoof et al., 2002).

As in the *NUDT21* CNV patient-derived lymphoblasts, *NUDT21* loss of function causes 3′ UTR shortening of *MECP2* in human neurons, as well as other genes strongly regulated by *NUDT21* in cancer-cell-line assays, such as *VMA21*, *LAMC1*, and *PAK1* (***Figure 4Cii & iii***) (Gennarino et al., 2015; Masamha et al., 2014). Also, similar to other cell-line assays, *NUDT21* loss of function causes widespread dysregulation of alternative polyadenylation, predominantly resulting in mRNA shortening (***Figure 4Ci-v & D***) (Brumbaugh et al., 2018; Gruber et al., 2012; Masamha et al., 2014). Of the misregulated genes, 230 have a gnomAD probability of loss-of-function intolerance over 0.9, 43 cause intellectual disability when mutated, and of those, 18 are autosomal or X-linked dominant (***Supplemental data 1***) (Lek et al., 2016; OMIM, 2019; Vissers et al., 2015). In addition to *MECP2*, misregulation of some of these genes is likely contributing to pathogenesis.

Notwithstanding disrupted RNA localization, the most apparent consequence of misregulated APA is altered protein levels (Tian and Manley, 2016). As expected, for many of the genes with significantly misregulated APA, we also see concordant changes in protein levels (***Figure 4E***). Most commonly, *NUDT21* loss results in shorter mRNAs and increased protein, such as with MeCP2 and VMA21, which respectively had protein-level increases of 50% and almost 300% (***Figure 4Cii-iii & E***). However, there are exceptions that further clarify the relationship between *NUDT21* loss and protein levels. Occasionally, the shortened mRNA will lose its stop codon and so likely undergo non-stop decay, resulting in reduced protein, such as with *ZCCHC6* (***Figure 4Civ & E***). Conversely, as with *KIF9*, *NUDT21* reduction occasionally induces synthesis of longer mRNAs, and this could paradoxically increase protein levels if the longer mRNAs gain a stop codon and cease to be degraded by non-stop decay (***Figure 4Cv & E***). Intriguingly, the degradation of KIF9 and others through non-stop decay in the control neurons shows an example of non-stop decay being used for normal gene regulation, rather than just as a quality control mechanism. We observed that ∼15% of the genes with altered APA have a change in the percentage of mRNA isoforms undergoing non-stop decay. However, the majority follow the trends seen in cancer cell lines: *NUDT21* loss of function in neurons causes widespread 3′ UTR shortening and increased protein levels (***Figure 4D & E***).

## DISCUSSION

Here we show that a partial loss of *Nudt21* function causes learning deficits and cortical hyperexcitability in mice. Further, we show that partial loss of *NUDT21* function broadly disrupts gene expression in human neurons via widespread misregulated alternative polyadenylation and protein levels. Alongside our previous discovery that patients with *NUDT21*-spanning deletions have intellectual disability and gnomAD data that *NUDT21* is a highly constrained gene, our results provide strong evidence that partial loss of *NUDT21* function causes intellectual disability (Gennarino et al., 2015; Lek et al., 2016).

Because a relatively small reduction in CFIm25 protein was sufficient to cause deficits, these data suggest that individuals with missense variants in *NUDT21* that affect its function may also have intellectual disability. Moreover, duplications of *NUDT21*, which we previously showed are associated with ID, should lead to comparable dysregulation of alternative polyadenylation and protein levels, including reduced MeCP2, and thus cause disease (Gennarino et al., 2015).

Beyond providing insight into *NUDT21*-associated disease, these data provide useful perspectives on the broader field of pediatric neurodevelopmental disease research. They illustrate the importance of protein-level homeostasis. A mere 30% reduction in CFIm25 protein was sufficient to cause learning deficits in mice, and mice are neurologically less sensitive to genetic insult than humans (Tan and Zoghbi, 2018). Further, many genes associated with neurodevelopmental diseases encode proteins that regulate or affect transcription (Rubeis et al., 2014; Vissers et al., 2015; Yin and Schaaf, 2017). *NUDT21* provides another layer to this family of genes insofar as it broadly affects alternative polyadenylation and protein levels. As a group, these genes demonstrate how a partial loss or gain of function can result in large effects, and further show that neurons are particularly sensitive to protein-level disequilibrium.

Lastly, while the observed 50% increase of MeCP2 protein likely contributes to the symptoms seen in the *NUDT21* CNV patients, the transcriptional dysregulation in neurons following *NUDT21* loss is so widespread that there are probably other CFIm25 targets that mediate the patients’ condition. Most likely, patient symptoms result from the additive effects of numerous misregulated genes, including the dozens that are ID associated. This then points to normalization of CFIm25 levels as the most viable molecular therapeutic strategy for treating this disease.

## METHODS

### Generation of *Nudt21*^+/-^ mice

The Baylor College of Medicine Institutional Animal Care and Use Committee approved all mouse care and manipulation (IACUC, protocol AN-1013). We generated C57BL/6J *Nudt21*^flox/flox^ mice by inserting loxP sites flanking exons 2 and 3 of *Nudt21* by homologous recombination in embryonic stem cells (Ozgene) (Weng et al., 2019). We then crossed male *Nudt21*^flox/flox^ mice with female C57BL/6J *Sox2*-Cre hemizygous mice to obtain *Nudt21*^+/-^ mice. All oocytes from *Sox2*-Cre hemizygous females have Cre and can induce cre/lox recombination, even those haploid oocytes that do not have the Cre transgene (Hayashi et al., 2002). We confirmed successful cre/lox recombination by Sanger sequencing. To eliminate potential confounding from recombination mosaicism, we first crossed the *Nudt21*^+/-^ F1 generation with wild-type C57BL/6J mice (Jackson Lab) before establishing mating pairs with F2s to generate experimental cohorts

### Immunofluorescence

We dissected then hemisected brains from three seven-week-old *Nudt21*^flox/flox^ C57BL/6J mice and drop-fixed their brains in 4% PFA overnight with gentle rocking at 4°C. After fixation, we cryoprotected the brains by gentle rocking in 30% sucrose solution in PBS at 4°C until the tissue sank. We then froze the brains in OCT, cryosected 40µm sections, and stored them at 4°C.

For immunolabeling, we first blocked free-floating sections in 5% normal goat serum, 0.5% Triton-X in PBS for one hour at room temperature. We then incubated the sections in CFIm25 primary antibody (1:50, Proteintech 10322-1-AP) in blocking buffer overnight at 4°C, followed by three washes and Alexa Fluor 488 (1:1000, Molecular Probes) secondary antibody incubation for two hours at room temperature. Lastly, we labeled the nuclei with DAPI (1:10,000) and mounted the sections with Vectashield. We used a Leica TCS SP5 confocal microscope for imaging and ImageJ Fiji for image processing. We confirmed the specificity of the CFIm25 antibody with si*NUDT21* knockdown in HeLa cells.

### RT-qPCR

*RNA extraction*: We dissected and hemisected the brains of five-week-old *Nudt21*^+/-^ mice and their wild-type littermates. We immediately flash froze them in liquid nitrogen and stored them temporarily at -80°C. We later homogenized the half brains in TRIzol Reagent (ThermoFisher Scientific) with the Polytron PT 10-35 GT (Kinematica). We then isolated the RNA by chloroform phase separation, precipitated it with 2-propanol, and washed it with 75% ethanol. We eluted the purified RNA in water.

We extracted RNA from shRNA-infected, human ESC-derived neurons in a 12-well plate: 4 wells non-silencing scrambled (RHS4348) and 8 wells sh*NUDT21* (V2LHS_253272 and V2LHS_197948) (Dharmacon). We lysed the neurons in the tissue-culture plate with TRIzol Reagent (ThermoFisher Scientific) and immediately transferred the lysate to microfuge tubes for trituration. We then isolated the RNA by chloroform phase separation, precipitation with 2-propanol, washing with 75% ethanol, and eluting in water as with the mice.

*RT-qPCR*: We synthesized first-strand cDNA with M-MLV reverse transcriptase (Life Technologies), and performed qPCR with PowerUp SYBR Green Master Mix (ThermoFisher Scientific) using the CFX96 Real-Time PCR Detection System (Bio-Rad). We designed exon-spanning primers using the UCSC genome browser mm10 and hg38 assemblies (http://genome.ucsc.edu/) (Kent et al., 2002), and Primer3 (http://bioinfo.ut.ee/primer3/)

(Koressaar and Remm, 2007; Untergasser et al., 2012). We performed all RT-qPCR reactions in triplicate and determined relative cDNA levels by *NUDT21* threshold cycle (Ct) normalized to *GAPDH* Ct using the delta Ct method: relative expression (RQ) = 2^-(*NUDT21* average Ct - *GAPDH* average Ct). We present the data from human neurons infected with shRNA clone V2LHS_253272 because we used it for the PAC-seq experiment. We analyzed the data by two-tailed, unpaired t-test, and present it as mean ± SEM. *, **, ***, and **** denote P<0.05, P<0.01, P<001, P<0.0001.

### Primers

*Nudt21*:

Forward: 5′- TACATCCAGCAGACCAAGCC - 3′

Reverse: 5′- AATCTGGCTGCAACAGAGCT - 3′

*Gapdh*:

Forward: 5′- GGCATTGCTCTCAATGACAA - 3′

Reverse: 5′- CCCTGTTGCTGTAGCCGTAT - 3′

*NUDT21*:

Forward: 5′- CTTCAAACTACCTGGTGGTG - 3′

Reverse: 5′- AAACTCCATCCTGACGACC - 3′

*GAPDH*:

Forward: 5′- CGACCACTTTGTCAAGCTCA - 3′

Reverse: 5′- TTACTCCTTGGAGGCCATGT - 3′

### Western blot

*Protein extraction*: For the mice, we dissected and hemisected the brains of five-week-old *Nudt21*^+/-^ mice and their wild-type littermates. We immediately flash froze them in liquid nitrogen and stored them temporarily at -80°C. We later homogenized the half brains with the Polytron PT 10-35 GT (Kinematica) in lysis buffer: 2% SDS in 100 mM Tris-HCl, pH 8.5, with protease and phosphatase inhibitors (ThermoFisher Scientific) and universal nuclease (Pierce). We incubated the lysates on ice for ten minutes, then rotated them for 20 minutes at room temperature. We spun the samples at top speed for 20 minutes to remove membrane, then quantified the protein levels with the Pierce Protein BCA Assay kit (ThermoFisher Scientific). For the ESC-derived neurons infected with shRNA targeting *NUDT21*, our extraction protocol was similar, except we lysed them directly in their 12-well plate and rocked them for 20 minutes at room temperature.

*Western blot*: We diluted the protein to 1 μg/μL in reducing buffer (LDS and sample reducing agent (ThermoFisher Scientific)) and ran 10 μg/sample. We imaged the membranes with the LI-COR Odyssey and analyzed the data with LI-COR Image Studio (LI-COR Biosciences), comparing H3-normalized CFIm25 levels. We confirmed the specificity of the CFIm25 antibody by sh*NUDT21* knockdown in HEK293T cells. We present the data from human neurons infected with shRNA clone V2LHS_253272 because we used it for the PAC-seq experiment. We analyzed the data by two-tailed, unpaired t-test, and present it as mean ± SEM. *, **, ***, and **** denote P<0.05, P<0.01, P<001, P<0.0001.

*Antibodies:*

CFIm25: NUDT21 (2203C3): sc81109 (Santa Cruz)

H3: Histone H3 (D1H2) XP (Cell Signaling Technology)

### Mouse husbandry and handling

The Baylor College of Medicine Institutional Animal Care and Use Committee approved all mouse care and manipulation (IACUC, protocol AN-1013). We housed the mice in an AAALAS-certified level three facility on a 14-hour light cycle. The mice had ad libitum access to standard chow and water. We weighed a cohort of mice weekly from 4-12 weeks and a different cohort at 30 weeks. We tested both male and females in all experiments and only present the weight data separately because we never otherwise detected a difference between them. We were blinded to the mice’s genotype during all handling.

Genotyping

We determined the mice’s genotypes by PCR amplification of tail lysates.

Primers:

*Nudt21*^null^

Forward: 5′- ACAGATTAGCTGTTAGTACAGG - 3′

Reverse: 5′- GAAGAACCAGAGGAAACGTGAG - 3′

Wild-type *Nudt21*

Forward: 5′- AGGAGGCTGACATGGATTGTT - 3′

Reverse: 5′- TCTTCTCCTGGGTTAAGTTCCC - 3′

### Behavioral tests

Because neurodevelopmental disease loss-of-function mouse models typically have more pronounced phenotypes when they are older, we started our behavioral battery on 30-week-old mice. We performed the assays on two cohorts. We tested the mice during their light cycle, typically between 11 AM and 5 PM. Prior to each assay, we habituated the mice to the testing facility for 30-60 minutes. The investigators were blind to the mice’s genotypes during all assays.

#### Conditioned fear

We first habituate the mice for 30 minutes in an adjacent room on each day of the test. On day one, we conditioned the mice by placing them in the habitest operant cage (Coulbourn) for a training session. The training consists of two minutes habituation, then a 30 second 85 dB tone followed by a foot shock of 1.0 mA for two seconds. After another two minutes, the 30 second 85 dB tone is played again (the training day cue). Throughout the experiment, except for the two seconds during the foot shock, the FreezeFrame3 system (Coulbourn/Actimetrics) recorded the mice’s movement and freezing episodes. On day two, we performed the contextual and cued fear assays. For contextual fear, we returned the mice to the test chambers precisely as we had done during the training, and recorded their freezing in the chamber for five minutes. We waited two hours before beginning the cued fear assay. We first changed the holding cages and test chamber shape, color, texture, scent, and lighting to make the experience as unrecognizable as possible to the mice. We then placed them in the modified chamber, and after three minutes played the 30 second 85 dB tone, then recorded their freezing for the following three minutes. We analyzed the difference in freezing between the two groups after the second sound cue (training cue) on day one, throughout the contextual fear test, and after the sound cue in the cued fear test, each by unpaired, two-tailed t-test. We excluded data for all the mice from one contextual fear trial. We suspect there was a technical error in the collection of those data: the FreezeFrame3 system recorded them as freezing far more than the mice in any other trial (50-80%) and two of the three were significant outliers in the Grubbs test.

#### Open field

We lit the room to 200 lux and set the ambient white noise to 60 dB during habituation and throughout the test. We placed each mouse in the open field, a 40 x 40 x 30 cm chamber equipped with photobeams (Accuscan Instruments), and recorded their activity for 30 minutes. We analyzed total distance and center tolerance (center distance/total distance) by unpaired, two-tailed t-test.

#### Morris water maze

Our Morris water maze experiment took pl ace in a 120 cm diameter pool of water. We hid a 10 cm X 10 cm platform 0.5-1 cm underwater in the Southeast quadrant. On each wall of the testing room, we taped brightly colored shapes that the mice can use for orientation. Our experiment spanned four days. Each day, we set the lighting to 60 lux and habituated the mice in the testing room in their home cages for 30 minutes, then ten minutes in holding cages. Prior to the first day’s experiment, we introduced the mice to the invisible platform by placing them on it for ten seconds. We next pulled the mice into the water and let them swim for ten seconds to ensure they could swim, then placed them directly in front of the platform to confirm they could climb back on to it. We tested the mice in two training blocks per day for the four days. Each training block consisted of four trials. For each trial, we placed the mice in a different quadrant of the pool (North, South, East, or West); the quadrant order was the same for every mouse in the trial, but different for every trial. We removed the mice after they found the platform, or if they did not find that platform, we guided them to the platform to rest on it for ten seconds before removing them. We used an EthoVision XT automated video tracking system (Noldus Information Technology) to track the mice’s location, speed, and latency to find the platform. After the second training block on the fourth day, we immediately performed the probe trial: we removed the platform and placed the mice in a new location, the Northwest quadrant, and tracked them for 60 seconds. We analyzed their speed by unpaired, two-tailed t-test; their latency to find the platform by two-way, repeated measures ANOVA (genotype*block); and their time in each quadrant in the probe trial by two-way, repeated-measures ANOVA (genotype*quadrant).

#### Animal behavior statistical analysis

From previous experience, we know a sample size of 14 animals is sufficient to detect meaningful phenotypic differences in a neurobehavioral battery (Chao et al., 2010; Lu et al., 2017; Samaco et al., 2012). We analyzed all the data with Prism 7 (Graphpad), following Graphpad’s recommendations. We used unpaired, two-tailed t-tests for all simple comparisons, and two-way repeated measures ANOVAs for all two-factor comparisons. We present all data as mean ± SEM. *, **, ***, and **** denote P<0.05, P<0.01, P<0.001, P<0.0001.

### Video electroencephalography (EEG) and spike counting

*Surgery and data recordings*: The Baylor College of Medicine Institutional Animal Care and Use Committee approved all research and animal care procedures. We tested eight *Nudt21*^+/-^ mice and six wild-type littermate controls. Experimenters were blind to the mouse genotype. We secured 54-week-old mice on a stereotaxic frame (David Kopf) under 1-2% isoflurane anesthesia. Each mouse was prepared under aseptic condition for the following recordings: the cortical EEG recording electrodes of Channels 1 and 2 were made of Teflon-coated silver wires (bare diameter 127 µm, A-M systems) and implanted in the subdural space of the parietal cortex and frontal cortex, respectively, with reference at the midline over the cerebellum. The electrode of the third channel, made of Teflon-coated tungsten wire (bare diameter 50 µm, A-M systems) was stereotaxically aimed at the hippocampal dentate gyrus (1.9 mm posterior, 1.7 mm lateral, and 1.8 mm below the bregma) with reference in the ipsilateral corpus callosum (Paxinos and Franklin, 2001). In addition, Teflon-coated silver wires were used to record the electromyogram (EMG) in the neck muscles to monitor mouse activity. All of the electrode wires together with the attached miniature connector sockets were fixed on the skull by dental cement. After two weeks of post-surgical recovery, mice received three two-hour sessions of EEG/EMG recordings over a week. Signals were amplified (100x) and filtered (bandpass, 0.1 Hz - 1 kHz) with the 1700 Differential AC Amplifier (A-M Systems), then digitized at two kHz and stored on disk for off-line analysis (DigiData 1440A and pClamp10, Molecular Devices). The time-locked mouse behavior was recorded by ANY-maze tracking system (Stoelting Co.).

*EEG data analysis*: Abnormal synchronous discharges were manually identified when the sharp positive deflections exceeding twice the baseline and lasting 25-100 ms (Roberson et al., 2011). We counted the number of abnormal spikes over the recording period using Clampfit 10 software (Molecular Devices, LLC) and averaged the spike numbers across sessions for each animal. Since the data follow a lognormal distribution, we statistically compared the genotypes with the Mann-Whitney test using Prism 7 (Graphpad). The measure of central tendency is the geometric mean and * indicates P<0.05.

### shRNA lentivirus production and titer assessment

#### Virus production

We made several viruses that express shRNAs targeting *NUDT21*. We transfected 45 ug DNA into 80%–90% confluent, low-passage HEK293T cells (ATCC CRL-3216; RRID:CVCL_0063) in 150 mm dishes at a 4:3:1 ratio of pGIPz, psPAX2, pMD2.G with TransIT-293 transfection reagent (Mirus, MIR 2706). The following day, we changed the media to 10 mL. At 48 and 72 hours, we collected and pooled their media, then centrifuged at 4000 x g for ten minutes and filtered the supernatant through a Poly-ethersulfone filter (VWR, 28145-505) to remove cellular debris. We concentrated the virus 100-fold with Lenti-X concentrator (Clontech, 631231) following the manufacturer’s recommendations before aliquoting and freezing at -80C.

#### Titer assessment

We measured the viral titer using Open Biosystems pGIPZ method (Thermo Fisher Scientific). We plated 5 × 10^4^ HEK293T cells in a 24-well plate. The following day, we made a serial dilution of the virus in a 96-well plate, and when the HEK293T cells reached ∼50% confluency, we infected them with the diluted virus. We cultured the cells for two days, then counted the tGFP colonies with Axiovert 25 microscope (Zeiss) microscope X-cite 120 lamp (ExFo) to determine the viral titer. Our viruses had 10^9^ transducing units/mL.

#### Confirmation

Compared to non-silencing scrambled control virus (RHS4348, Dharmacon), we confirmed *NUDT21* knockdown efficacy in HEK293T cells (ATCC CRL-3216; RRID:CVCL_0063), and selected the two most efficient shRNAs for our studies: V2LHS_197948 and V2LHS_253272 (Dharmacon).

### Human embryonic stem cell (hESC)-derived neuron culture

We used WA09 (H9; RRID:CVCL_9773) female embryonic stem cells (ESCs) to generate human neurons as previously described (Jiang et al., 2019). We differentiated neural progenitors into human neurons over three weeks, changing the media every 3 d. Afterwards, we passaged the neurons with trypsin. Three days after passaging, we infected the neurons with lentiviruses containing pGIPZ shRNA clones at a multiplicity of infection of 10. We verified the tropism and infectivity of the virus using the tGFP reporter signal. At day 3 after infection, we treated the neurons with puromycin (0.75–1.25 g/ml) for 6 days to select for infected cells. We cultured the cells for 60 days after infection, changing the media three times per week. We then aspirated all the media and washed the cells with PBS before freezing them at -80C for later RNA and protein extraction.

### Poly(A) click-seq

#### RNA extraction

We extracted RNA from shRNA-infected, human ESC-derived neurons in a 12-well plate: 4 wells non-silencing scrambled (RHS4348) and 8 wells sh*NUDT21* (V2LHS_253272 and V2LHS_197948) (Dharmacon). We know from past experience that we need at least three samples for RNA- and PAC-seq analysis (Elrod et al., 2019; Maio et al., 2018; Routh et al., 2017; Tan et al., 2016). We prepared four samples per genotype to allow for loss of one sample. We lysed the neurons in the tissue-culture plate with TRIzol Reagent (ThermoFisher Scientific) and immediately transferred the lysate to microfuge tubes for trituration. We then isolated the RNA by chloroform phase separation, precipitation with 2-propanol, washing with 75% ethanol, and eluting in water.

#### Library preparation and sequencing

We prepared sequencing libraries as previously described (Routh et al., 2017). We reverse transcribed 1 ug of total RNA with the partial P7 adapter (Illumina_4N_21T) and dNTPs with the addition of spiked-in azido-nucleotides (AzVTPs) at 5:1. We click-ligated the p5 adapter (IDT) to the 5′ end of the cDNA with CuAAC. We then amplified the cDNA for 21 cycles with Universal primer and 3′ indexing primer and purified it on a 2% agarose gel by extracting amplicon from 200-300 base pairs. We pooled the libraries and sequenced single-end, 75 base-pair reads on a Nextseq 550 (Illumina).

P7 adapter (Illumina_4N_21T): GTGACTGGAGTTCAGACGTGTGCTCTTCCGATCTNNNNTTTTTTTTTTTTTTTTTTTTT)

P5 adapter (IDT): 5′HexynylNNNNAGATCGGAAGAGCGTCGTGTAGGGAAAGAGTGTAGATCTCGGTGGT CGCCGTATCATT

Universal primer: AATGATACGGCGACCACCGAG

Example 3′ indexing primer: CAAGCAGAAGACGGCATACGAGATCGTGATGTGACTGGAGTTCAGACGTGT

### PAC-seq data analysis

We sequenced each library to a depth of ∼20 million, 75 base pair (bp), single-end reads. For each sample, we obtained raw reads in a zipped fastq format. We used fastp for initial quality control (Chen et al., 2018). We filtered adapter contamination (AGATCGGAAGAGC) using the -a option. We trimmed the first 6 nucleotides and any PolyG tail nucleotides with the -f and -g options. And removed reads shorter than 40 nucleotides (nt) after trimming, using the -l option.

#### Read alignment

We downloaded raw FASTA sequences and annotations of the Human genome build GRCh38 from the UCSC table browser tool (Karolchik et al., 2004), and aligned trimmed reads to the reference genome with Bowtie 2 version 2.2.6 with parameters: -D 20 -R 3 -N 0 -L 20 -i S,1,0.50 (very-sensitive-local) (Langmead and Salzberg, 2012). We indexed the reference genome using bowtie2-build default settings, and saved sample-wise alignments as Sequence Alignment Map (SAM) files. We then used SAMtools V0.1.19 “*view”*, “*sort*” and “*index*” modules to convert SAM files to Binary Alignment Maps (BAM), coordinate sort, and index (Li et al., 2009).

#### Sample quality

We obtained gene level counts as the sum of total reads mapped to their respective genes. shRNA clone V2LHS_253272 gave more consistent *NUDT21* knockdown, so we used those samples in our analysis. We sequenced four samples from both the control and *NUDT21* knockdown conditions, but excluded one control sample with <10 million reads, and one knockdown sample with unusually low *NUDT21* levels (<2% of other knockdown samples). We then performed principle component analysis (PCA) on gene counts as previously described and confirmed that the treatment groups clustered (***Figure 4—figure supplement 1A***) (Yalamanchili et al., 2017).

#### Alternative poly-adenylation (APA) analysis

We computed strand-specific coverage of the features (mRNA cleavage sites) from BAM files using bedtools v2.25.0 (Quinlan and Hall, 2010) “*genomecov”* module. We pooled sample-wise feature files to get a comprehensive list of polyadenylation (p(A)) sites across all the samples.

We merged overlapping features and features within 15 nucleotides using the bedtools “*merge*” module. We mapped features to genes if they were within the gene body or within 16 kilobases (kb) downstream of the annotated transcription end site and do not overlap with any other gene, which is far enough to detect the longest known 3′UTRs (Miura et al., 2013; Quinlan and Hall, 2010). We then mapped these features to known, annotated human polyadenylation sites downloaded from PolyA_DB3 (Wang et al., 2017). Because the annotations were in the hg19 coordinate system, we converted them to the GRCh38 coordinate system with the liftOver program from the UCSC genome browser portal. With a read length of 75 nucleotides, we may not have reached the actual cleavage sites. Thus, each annotated polyadenylation site was considered a 70-nucleotide feature for mapping our features to annotated p(A) sites. Our features that did not overlap with at least 1 bp of any of the annotated p(A) sites were considered novel p(A) sites.

For calling novel p(A) sites, we used a sliding window approach to filter out any potential misprimed sites, i.e. potential cleavage sites where the poly-d(T) reverse transcription primer bound genomic adenines in the body of the mRNA rather than the poly(A) tail. We filtered out features with more than 12 genomic adenines in a 20 bp window 10 nt up- and 50 nt downstream of the potential cleavage site.

We quantified resultant novel and annotated p(A) sites across all samples using featureCounts (Liao et al., 2013). To minimize potential bias from under-covered p(A) sites, we retained only features with at least 5 reads in two samples and 1 read in the remaining sample in either of the conditions. We also filtered out p(A) sites accounting for less than 10% of the reads mapped to a gene in both the control and *NUDT21* knockdown conditions. We considered genes with mean read counts <5 in either condition not expressed and so excluded them.

Because about half of genes that undergo APA have greater than two p(A) sites, we wanted to use a statistic that could account for changes in all of the p(A) sites (Derti et al., 2012). Thus, we identified changes in alternative polyadenylation site usage using the Dirichlet multinomial test in the DRIMSeq package, and computed adjusted p-values as previously described (Nowicka and Robinson, 2016; Zhao et al., 2013). We quantified the magnitude and direction of change in polyadenylation site usage as an mRNA lengthening score. We summed the weighted read counts per million (cpm) from each p(A) site in a gene to get that gene’s mRNA length score.

We assigned the weights in decreasing order from the most distal p(A) site as below:

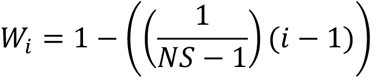

where *W_i_*is the weight of *i^th^* distal polyadenylation site of a gene and *NS* is the of total number of polyadenylation sites mapped to it. Thus, in a gene with 3 p(A) sites, the cpm at the most promoter-distal site would all be counted (weight of 1) in the length score, the cpm at the intermediate site would be weighted by 0.5, and the cpm at the proximal site would not be counted (weight of 0). We then averaged the scores across samples and took the difference of treated to controls for the relative mRNA length change.

#### Disease association determination

For the probability of loss of function intolerance (pLI) of all the genes, we used gnomAD (Lek et al., 2016). In addition, we cross-referenced the misregulated genes with published genetic studies to see which were known to cause intellectual disability when mutated, then looked to the Online Mendelian Inheritance in Man (OMIM) database to see if the pathological variants were dominant or recessive (OMIM, 2019; Vissers et al., 2015).

### Mass spectrometry

We processed, measured, and analyzed the sample as previously described (Saltzman et al., 2018). We summarize the main steps below.

#### Lysis and Digestion

We pelleted and lysed the shRNA-infected human neurons with three freeze (LN2) and thaw (42°C) cycles in 50 μL of ammonium bicarbonate + 1mM CaCl_2_. We then boiled the lysate at 95°C for two minutes with vortexing at 20 second intervals, and then centrifugation at 21,000 rcf. We digested 50 μg of total protein with a 1:20 solution of 1 μg/μL trypsin:protein overnight at 37°C with shaking and then again with a 1:100 solution of 1 μg/μL trypsin:protein for 4 hours. We extracted the peptides with 80% acetonitrile + 0.1% formic acid solution, followed by centrifugation at 10,000 g, and vacuum drying.

#### Off-Line Basic pH Reverse Phase Peptide Fractionation

A “15F5R” protocol was used for off-line fractionation as described before(Saltzman et al., 2018). We filled a 200 μl pipette tip with 6 mg of C18 matrix (Reprosil-Pur Basic C18, 3 μm, Dr. Maisch GmbH) on top of a C18 disk plug (EmporeTM C18, 3M). We dissolved 50 μg of vacuum-dried peptides with 150 μl of pH10 ABC buffer and loaded it onto the pre-equilibrated C18 tip. We eluted bound peptides with fifteen 2%-step 2-30% gradient of ACN non-contiguously combined into five pools, and then vacuum dried.

#### Mass Spectrometry

We analyzed fractionated peptides on an Orbitrap Fusion mass spectrometer coupled with the Nanospray Flex ion sources and an UltiMate 3000 UPHLC (Thermo Fisher Scientific). For each run, we loaded approximately one microgram of peptide onto a two cm 100 μm ID pre-column and resolved it on a twelve cm 100 μm ID column, both packed with sub-twi μm C18 beads (Reprosil-Pur Basic C18, Catalog #r119.b9.0003, Dr. Maisch GmbH). We maintained a constant flow rate for 100-minute 2-28% B gradient elutions, where A is water and B is 90% acetonitrile, both with 0.1% formic acid.

#### Proteome Discoverer (Mascot-based) Search and Protein Inference/Quantification

We used the Proteome Discoverer software suite (PD version 2.0.0.802; Thermo Fisher Scientific) to search the raw files with the Mascot search engine (v2.5.1, Matrix Science), validate peptides with Percolator (v2.05), and provide MS1 quantification through Area Detector Module (Käll et al., 2007; Perkins et al., 1999). We matched MS1 precursors in a 350-10,000 mass range against the tryptic RefProtDB database digest (2015-06-10 download) with Mascot permitting up to 2 missed cleavage sites (without cleavage before P), a precursor mass tolerance of 20 ppm, and a fragment mass tolerance of 0.5 Da. We allowed the following dynamic modifications: Acetyl (Protein N-term), Oxidation (M), Carbamidomethyl (C), DeStreak (C), and Deamidated (NQ). For the Percolator module, we set the target strict and relaxed FDRs for PSMs at 0.01 and 0.05 (1% and 5%), respectively. We used gpGrouper (v1.0.040) for gene product inference and label-free iBAQ quantification with shared peptide distribution.

#### Data Normalization

Because *NUDT21* inhibition consistently increases protein levels across the proteome, normalizing to total protein levels in each sample would introduce bias. Instead, we normalized each sample to a group of ∼20 consistently expressed genes that we know are not directly regulated by *NUDT21* from the PAC-seq data.

#### Analysis

We performed a PCA as previously described to confirm that the treatment groups separated (***Figure 4—figure supplement 1B***) (Yalamanchili et al., 2017). We then computed protein level changes as log_2_ fold change values and plotted them against their relative mRNA length change (***Figure 4E***).

## Data availability

The PAC-seq data are available in the NCBI Gene Expression Omnibus, accession number GSE135384. The analysis code can be found at http://liuzlab.org/iNeuron_PACSeq.zip. The mass spectrometry proteomics data have been deposited to the ProteomeXchange Consortium via the PRIDE partner repository with the dataset identifier PXD014842 (Perez-Riverol et al., 2018).

## ETHICS

For animal experimentation, we housed up to five mice per cage on a 12-hour light cycle in a level 3, AALAS-certified facility and provided water and standard rodent chow ad libitum. The Institutional Animal Care and Use Committee for Baylor College of Medicine and Affiliates approved all procedures carried out in mice under protocol AN-1013.

## ACKOWLEDGEMENTS

We would like to thank the Zoghbi and Wagner lab members for critical feedback, particularly Nathan Elrod for his suggestions on analyzing the PAC-seq data. We would also like to thank the cores that provided services for the project: the Microscopy, Neuroconnectivity, Animal Behavior, and Human Neuronal Differentiation cores from the Jan and Dan Duncan Neurological Research Institute and BCM Intellectual and Developmental Disabilities Research Center (NIH U54 HD083092 from the Eunice Kennedy Shriver National Institute of Child Health and Human Development); the Next Generation Sequencing core at the University of Texas Medical Branch; and BCM Mass Spectrometry Core (supported by NIH P30 CA125123 and CPRIT RP170005). Lastly, we would like to thank our funders who made the work possible: the NIH National Institute of Neurological Disorders and Stroke—F30NS095449 (CEA), the Howard Hughes Medical Institute (HYZ), and the Intellectual and Developmental Disabilities Research Center—NIH U54 HD083092 from the Eunice Kennedy Shriver National Institute of Child Health and Human Development (HYZ). The content is solely the responsibility of the authors and does not represent the official views of the Eunice Kennedy Shriver National Institute of Child Health and Human Development, the National Institutes of Health, or the Howard Hughes Medical Institute.

